# PXGS: a Poly-Transgene Expression System based on Mutually Exclusive Splicing of *Dscam*

**DOI:** 10.1101/2024.10.27.620485

**Authors:** Renee Yin Yu, Alyeri Bucio-Méndez, Brian E. Chen

## Abstract

Biologists often need to investigate multiple genes simultaneously in an organism. However, it is currently not possible to express more than a few transgenes in an animal under conditional control. Here, we developed a technique based on the mutually exclusive splicing of the *Down Syndrome Cell Adhesion Molecule1* (*Dscam1*) gene in *Drosophila melanogaster* to achieve simultaneous transgene expression of 12 genes at a time. We show that the hypervariable *Dscam1* exon 4 region maintains its alternative splicing when placed in a UAS expression vector. Each of the twelve exon 4 alternates can be replaced with an exogenous gene of at least 10 kilobases and will express properly *in vivo* all under conditional genetic control. We demonstrate the expression of four different fluorophores placed in different exon 4 alternate positions in neural and non-neural cells *in vivo*. We validated the technique by rewiring *Drosophila* sensory neuron axons *in vivo* by simultaneously expressing several cell surface receptors within the neuron. This technology will also enable *Drosophila melanogaster* as a model system for synthetic biology research.

## Introduction

Proteins evolve within complexes of functional units, and in structural biology the detection of a protein’s structure within its multiprotein complex is important for understanding its biological function and evolutionary relationship (Ban et al., 2000; Bhattacharya, 2009; Stahl et al., 2024; Wiederstein et al., 2014). In cellular and systems biology, multiple molecules often need to be simultaneously manipulated to examine their roles and interactions within a cell or organism. In bacteria and yeast, multiple genes can be expressed using a promoter for each gene, but repeats of the same promoter can result in homologous recombination (Agmon et al., 2009). In multi-cellular organisms such as animals, it can be challenging to express multiple transgenes.

Multiple genes can be expressed using a single polycistronic mRNA strand (Kozak, 1999). For example, an internal ribosome entry site (IRES) is used by the poliovirus to initiate translation (Pelletier and Sonenberg, 1988), but can be used for bi-cistronic expression of two genes. An IRES sequence can be placed between an upstream gene and a downstream gene so that a ribosome can bind not only to the 5’ cap of the mRNA to translate the upstream gene, but also the IRES site to translate the downstream gene. However, the gene downstream of the IRES site is often translated at a much lower rate than the upstream gene and only two genes at a time is possible (Mizuguchi et al., 2000). Polycistronic expression can also be achieved using RNA sequences that encode cis-acting hydrolase element (CHYSEL) peptides (Doronina et al., 2008). CHYSEL polypeptides are commonly known as “2A” and “2A-like” peptides, and are used by RNA viruses to express each of its viral genes in a single mRNA strand. Synthesis of the CHYSEL polypeptide causes steric hindrance at the exit tunnel of the ribosome, causing the ribosome to skip the last bond of the CHYSEL peptide chain and restart on the next peptide (Donnelly et al., 1997; Martin D. Ryan GL, 2002; Ryan et al., 1991). Genes placed upstream and downstream of specific CHYSEL peptide sequences can be co-translationally co-expressed at an almost stoichiometric ratio (Lo et al., 2015). Up to four genes can be simultaneously expressed (quad-cistronic) using this technique, but at decreasing efficiency for each downstream gene (Liu et al., 2017). Finally, the MultiLabel technique in mammalian cells can express up to five genes (Kriz et al., 2010), and is based on the MultiBAC baculovirus expression system used in moth *Spodoptera frugiperda* cell lines (Bieniossek et al., 2012; Sari et al., 2016).

Alternative splicing is nature’s way of producing molecular diversity from a single gene. An extreme example of alternative splicing is the *Drosophila Down syndrome cell adhesion molecule1 (Dscam1)* gene. Dscam is a cell surface receptor that is involved in several aspects of neural development, neural plasticity, and immune recognition (Schmucker and Chen, 2009). The *Dscam1* gene can produce tens of thousands of different isoforms through mutually exclusive splicing. It has 20 constitutive exons and 4 variable exon clusters: 4, 6, 9, and 17 which can be alternatively spliced in a mutually exclusive manner within each cluster (Schmucker et al., 2000) (**Figure 1a**). Within each of these variable exon clusters, the cell will select one and only one of the possible alternatives to splice into the mRNA during each transcription event. In variable exon 4, there will be only 1 out of 12 alternates spliced, in variable exon 6 only 1 out of 48 will be chosen, in exon 9 only 1 out of 33 will be chosen, and in exon 17 one or the other alternate is chosen, all incorporated into a single mRNA strand (**Figure 1a**). This splicing occurs for every round of *Dscam1* transcription, so there will be many mRNA molecules that each contain a different exon variant at each of the variable exon clusters.

**Figure 1.**
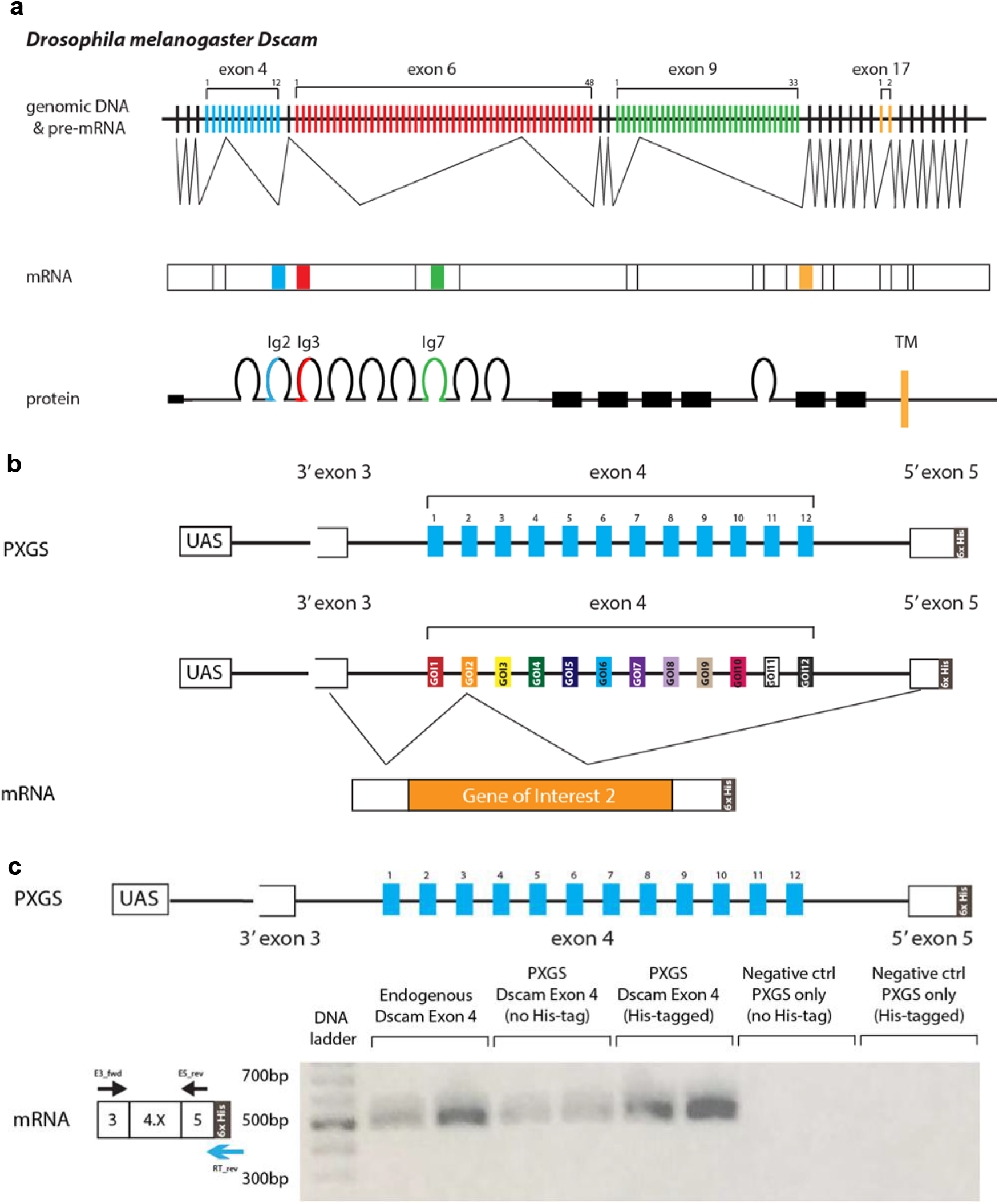
PXGS takes advantage of the mutually exclusive splicing property of *Dscam*. **a**, Endogenous *Dscam1* mRNA is mutually exclusively spliced in four exons during transcription, exons 4, 6, 9, and 17. For each of the mutually exclusively spliced exons, with every round of transcription, one and only one of the possible exon variants is in the final mRNA product and ultimately the protein product. **b**, The entire *Dscam* exon 4 cluster is inserted downstream of a *UAS* sequence to create the *PXGS* vector. A 6× histidine tag is added at the end of exon 5 to distinguish *PXGS* expression from endogenous *Dscam* exon 4 expression. Any gene of interest (GOI) can then be inserted to replace any of the 12 exon 4 alternates for expression. With each round of transcription, only a single transgene is selected for incorporation into the final mRNA product. **c**, Exon 4 mutually exclusive splicing is maintained in PXGS. *PXGS* vectors with and without a 6× histidine tag were co-transfected with *Actin5c-*Gal4 in *Drosophila S2* cells. As a positive control, reverse transcription (RT) of Exon 4 was performed on untransfected *S2* cells to identify the endogenous *Dscam* Exon 4 mRNA. RT using PXGS-specific reverse primers on *S2* cells was performed, with the no *Actin5c-*Gal4 co-transfection (PXGS only) as negative controls. PCR of Exon 4 in the PXGS vectors with and without a 6×His-tag showed DNA bands at the expected 500 bp to 600 bp sizes.

Here, we show that placing the entire *Dscam* exon 4 alternatively spliced cassette section into a DNA plasmid under UAS control, and replacing each of the twelve exon 4 variants with a gene, allows each gene to be expressed simultaneously in a cell. Cell-or tissue-specific expression is controlled using the Gal4/UAS system (Brand and Perrimon, 1993). We also show that alternative splicing of polycistronic dsRNA can induce RNAi *in vivo*. We validate our technology by rewiring *Drosophila* sensory neurons axons *in vivo* by simultaneously expressing several large cell surface receptors solely within the neuron. We call this new technology the poly-transgene expression system (PXGS).

## Results

### *Dscam* mutually exclusive splicing is conserved in the PXGS system

We first sought to verify that *Dscam* variable exon 4 would be properly spliced within an exogenous DNA construct. We extracted the last 300 bases of *Dscam* exon 3 through variable exon 4 to the first 30 bases of exon 5 from *Drosophila melanogaster* genomic DNA. A *6× Histidine* DNA sequence was added to the end of the exon 5 sequence as a genetic tag. We also wanted to test whether *Dscam* alternative splicing would be preserved after transcription under a UAS promoter (**Figure 1b**), to allow it to be expressed conditionally using a Gal4 driver (Brand and Perrimon, 1993). We inserted the 6.7 kilobasepair *Dscam* exon 4 fragment into the expression vector *pJFRC7-20XUAS*, and co-transfected it along with *Actin5C-Gal4* into *Drosophila melanogaster S2* cells (**Figure 1c**). To determine whether the endogenous exon 4 alternates were properly spliced, we performed reverse transcriptase PCR (RT-PCR) using the *6× Histidine* genetic tag to distinguish the UAS-transcripts from the endogenous *Dscam* mRNA transcripts. We found that in *S2* cells transfected with *UAS-DscamExon4*, the exogenous Exon 4 variants were produced at the expected sizes as the endogenous Exon 4 alternates (**Figure 1c**).

### Inserting fluorescent protein genes in the endogenous exon 4 alternates

The mutually exclusive splicing of *Dscam* exon 4 is determined by its intronic structures (Hong et al., 2021; Kreahling and Graveley, 2005; Xu et al., 2019; Yue et al., 2016), but it is not certain that the exonic 4 sequences do not contribute. First, we sought to insert the *green fluorescent protein* (*GFP*) transgene inside of (i.e., surrounded by) the endogenous exon 4 alternates, which would result in the endogenous sequences being treated as untranslated regions (UTRs) of mRNA flanking the transgene. Our goal was to then systematically narrow down the minimum endogenous *Dscam* exonic 4 sequences required for proper splicing. In parallel, we tried to completely replace the entire exon 4.1 alternate with the *GFP* gene. To our surprise, the *GFP* mRNA and functional fluorescence were expressed in *S2* cells when the entire exon 4.1 was replaced (**Supplemental Figure 1**). Thus, we found that only the intronic sequences within variable exon 4 are required for proper mutually exclusive splicing.

### Replacing exon 4 variants with fluorophores

We next sought to verify that other exon 4 variants could be completely replaced to express any gene of interest within our *UAS-DscamExon4* construct. We replaced alternate exons 4.1, 4.2, 4.11, and 4.12 with *GFP, blue fluorescent protein* with a nucleolar localization signal (*BFP*_*nols*_), *near-infrared fluorescent protein* (*iRFP*), and *red fluorescent protein* (*RFP*) transgenes, respectively. After verification of their expression in *S2* cells using RT-PCR, we generated a transgenic fly containing the *UAS-DscamExon4* construct, *UAS-DscamExon4_GFP*^*4*.*1*^*-BFP*_*nols*_^*4*.*2*^*-iRFP*^*4*.*11*^*-RFP*^*4*.*12*^. We crossed this fly to the pan-neuronal Gal4 driver *nSyb-Gal4*, and found that all four fluorophores were weakly to moderately expressed but not in all neurons (**Figure 2**). If Exon 4 alternative splicing is random, then after multiple rounds of transcription eventually all colors should be expressed. If Exon 4 splicing is deterministic, then some neurons may never express the splicing factors that select the missing colors. In other words, some neurons may not express specific splicing factors to allow for expression of specific Exon 4 alternates. An important caveat for this *UAS-DscamExon4* construct is that for each round of transcription driven by the nSyb-Gal4, a neuron may splice any of the eight Exon 4 endogenous alternates (i.e., producing non-fluorescence) or any of the four fluorophores, so that the “signal” is diluted and divided, respectively, compared to if nSyb-Gal4 were driving expression of a standard UAS-fluorescent protein construct.

**Figure 2.**
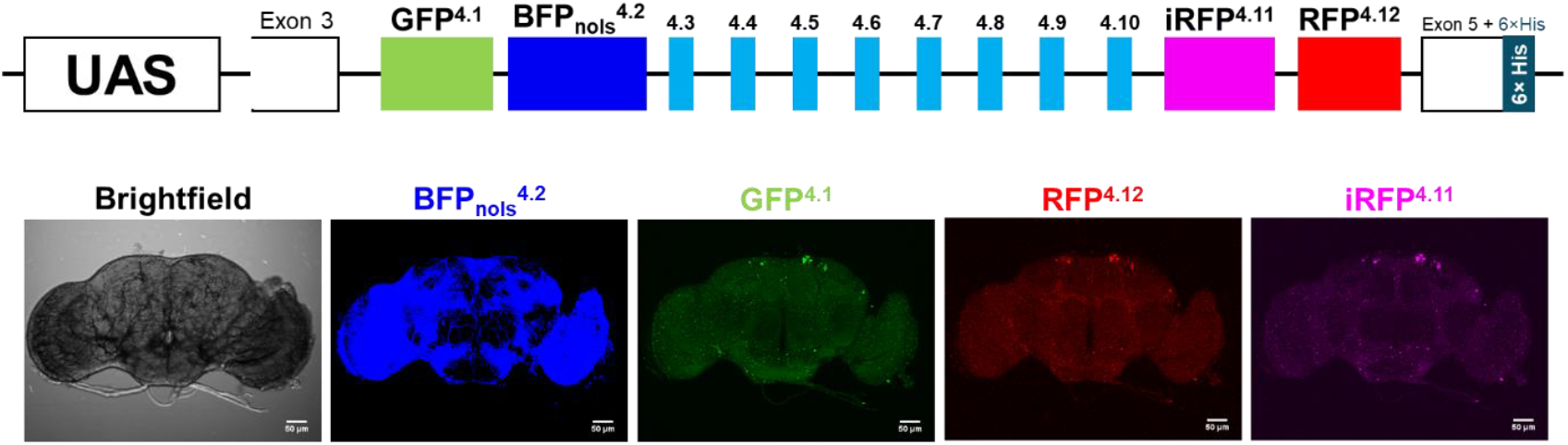
PXGS can express multiple fluorophores simultaneously *in vivo*. Crossing the pan-neuronal *nSyb-Gal4* driver line with the transgenic fly *UAS-PXGS_GFP*^*4*.*1*^*-BFP*_*nols*_^*4*.*2*^*-iRFP*^*4*.*11*^*-RFP*^*4*.*12*^ demonstrates that nearly all neurons can splice and express multiple transgenes *in vivo*. However, the fluorescence intensity for any single color was low as its production was divided among 12 Exon 4 alternates.

Regardless if *Dscam* Exon 4 is random or deterministic, we show that Exon 4 alternates can be replaced by a gene of interest for poly-transgene expression and can be conditionally controlled by Gal4 expression. After transcription and splicing, any extraneous DNA surrounding the transgene such as Exon 3, Exon 5, and the *6× Histidine* tag are untranslated mRNA regions (UTRs) and are not synthesized by the ribosome, as each transgene contains their own start and stop codons.

### PXGS can drive expression in non-neuronal cells

*Dscam* is expressed in all cells, based on gene expression atlases for *Drosophila*, albeit at varying levels (Corrales et al., 2022; Li et al., 2022). *Dscam* has traditionally been studied in the nervous and immune systems. To verify that our new poly-transgene expression system (PXGS) can be used in cells beyond neurons and immune cells, we generated a multi-color PXGS reagent to label any cell that expresses *Dscam* by replacing ten exon 4 alternates with fluorophores. We differentially localized GFP, BFP, RFP, and iRFP to the membrane (mCD8::GFP), the nucleus (nols), or the mitochondria (COX8::RFP), and generated the construct *PXGS_iRFP*_*nols*_^*4*.*1*^*-COX8::RFP*^*4*.*2*^*-BFP*_*nols*_^*4*.*3*^*-mCD8::GFP*^*4*.*4*^*-iRFP*_*nols*_^*4*.*5*^*-COX8::RFP*^*4*.*6*^*-mCD8::GFP*^*4*.*8*^*-BFP*_*nols*_^*4*.*9*^*-iRFP*_*nols*_^*4*.*10*^*-BFP*_*nols*_^*4*.*12*^. We verified the PXGS expression in *S2* cells by co-transfection with *Actin5C-Gal4* (**Supplemental Figure 2a**) and proceeded to generate transgenic flies. We crossed the *UAS-PXGS_iRFP*_*nols*_^*4*.*1*^*-COX8::RFP*^*4*.*2*^*-BFP*_*nols*_^*4*.*3*^*-mCD8::GFP*^*4*.*4*^*-iRFP*_*nols*_^*4*.*5*^*-COX8::RFP*^*4*.*6*^*-mCD8::GFP*^*4*.*8*^*-BFP*_*nols*_^*4*.*9*^*-iRFP*_*nols*_^*4*.*10*^*-BFP*_*nols*_^*4*.*12*^ flies to the pan-neuronal *nSyb-Gal4* and imaged the brains to verify expression *in vivo* (**Supplemental Figure 2b**). Neurons were strongly fluorescent with green at the cell membrane, intracellular punctate red mitochondria, and overlapping blue and infrared nuclei. However, the blue channel was often dominated by autofluorescence of the tracheal tubes.

Crossing the *UAS-PXGS_iRFP*_*nols*_^*4*.*1*^*-COX8::RFP*^*4*.*2*^*-BFP*_*nols*_^*4*.*3*^*-mCD8::GFP*^*4*.*4*^*-iRFP*_*nols*_^*4*.*5*^*-COX8::RFP*^*4*.*6*^*-mCD8::GFP*^*4*.*8*^*-BFP*_*nols*_^*4*.*9*^*-iRFP*_*nols*_^*4*.*10*^*-BFP*_*nols*_^*4*.*12*^ flies to the pan-glial *Repo-Gal4* driver revealed expression of all four fluorophores across the brain (**Figure 3**). We crossed these PXGS flies to *Tubulin-Gal4* for ubiquitous expression across the animal. Fluorescence expression was highest in the brain, but we also observed fluorescence in muscle. Multi-nucleated muscle cells were labelled with blue and near-infrared fluorescence, green cell membrane, and red mitochondria puncta (**Figure 3**). Similar to previous studies, we observed the mCD8::GFP expression concentrated at the immediate area surrounding the nucleus (Ralston and Hall, 1989).

**Figure 3.**
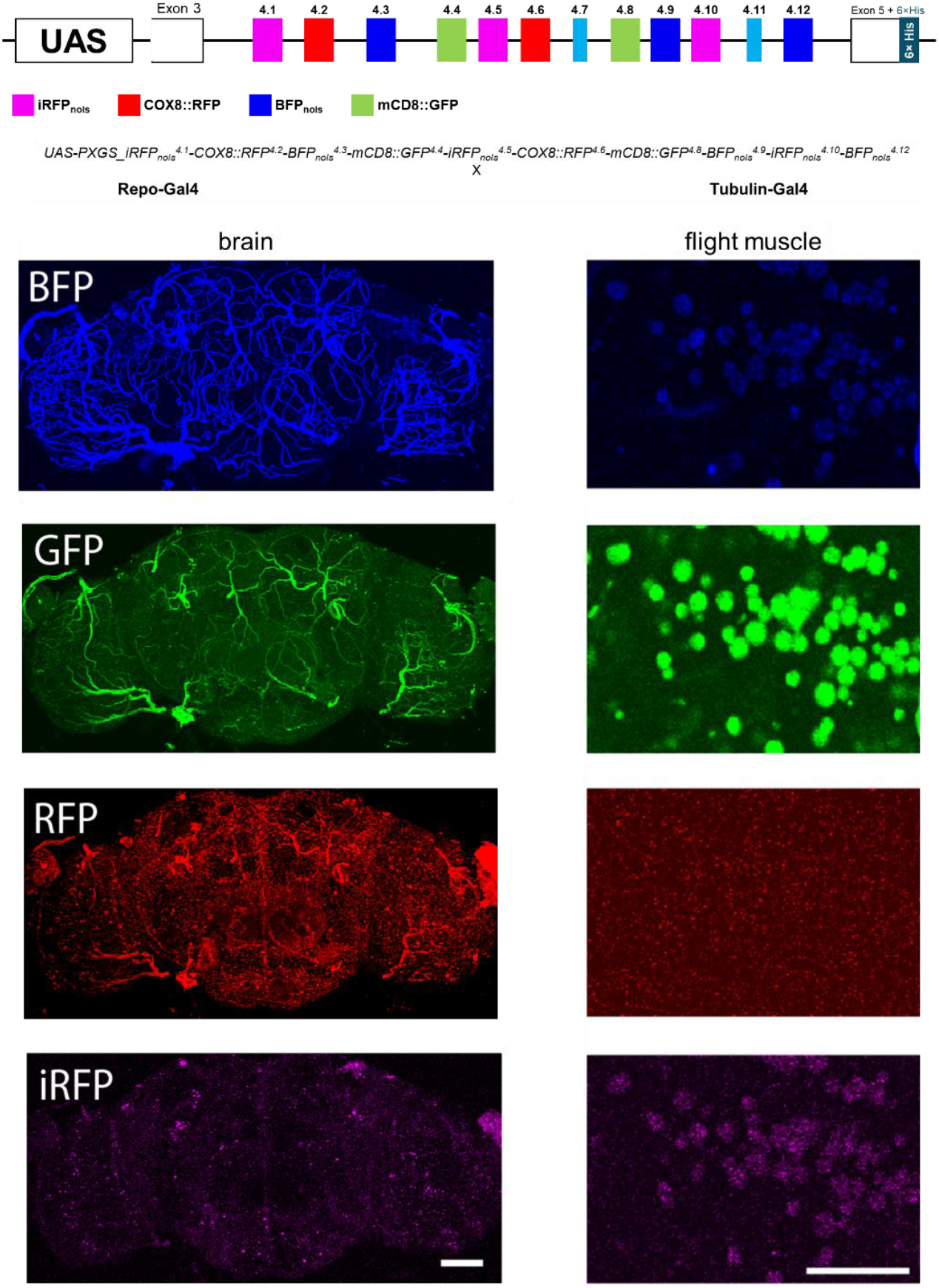
PXGS can drive expression in non-neural cells. Crossing the transgenic fly *UAS-PXGS_iRFP*_*nols*_^*4*.*1*^*-COX8::RFP*^*4*.*2*^*-BFP*_*nols*_^*4*.*3*^*-mCD8::GFP*^*4*.*4*^*-iRFP*_*nols*_^*4*.*5*^*-COX8::RFP*^*4*.*6*^*-mCD8::GFP*^*4*.*8*^*-BFP*_*nols*_^*4*.*9*^*-iRFP*_*nols*_^*4*.*10*^*-BFP*_*nols*_^*4*.*12*^ to the glial *Repo-Gal4* driver (left column) or to the ubiquitous *Tubulin-Gal4* driver (right column) showed expression in non-neural cells. The subcellular localization of the fluorophores was observed in the flight muscle (right column), but was less apparent in glial cells in the brain (left column). Scale bars are 50µm.

### Functional expression of PXGS transgenes

The genes for fluorescent proteins are ≤1 kilobase in size. To demonstrate that PXGS can express large, functionally relevant genes, we chose eight cell surface receptors to mis-express in a mechanosensory neuron. The eight genes were *Bsg, dpr8, dpr12, kek1, kirre, sli, Toll-6*, and *tutl*. We created three PXGS constructs: *UAS-PXGS_dpr8*^*4*.*1*^*-dpr12*^*4*.*2*^*-Gal4*^*4*.*3*^, *UAS-PXGS_kek1*^*4*.*7*^*-kirre*^*4*.*8*^*-tutl*^*4*.*9*^, and *UAS-PXGS_ Toll-6*^*4*.*10*^*-Bsg*^*4*.*11*^*_sli*^*4*.*12*^ and verified their mRNA expression in *S2* cells (**Supplemental Figure 3**) before generating fly lines. The gene for Gal4 itself was inserted into the Exon 4.3 position to create a transcriptional positive feedback loop for continuous gene expression. We used the *455-Gal4* driver line to mis-express these cell surface receptors solely within the pSc mechanosensory neuron (Cvetkovska et al., 2013; Dos Santos et al., 2019; Kays et al., 2014; Neufeld et al., 2011). The pSc mechanosensory neuron has a stereotyped axonal targeting pattern within the central nervous system, which can be used as an assay to identify molecules involved in axonal growth and targeting. *Dscam* isoforms are required for proper neural wiring (Chen et al., 2006), so overexpression of the PXGS constructs may interfere with the alternative splicing of endogenous *Dscam* isoforms by competing for the same splice factors, in a “sponging” effect. To account for this, we crossed the *455-Gal4* driver to the *UAS-PXGS_iRFP*_*nols*_^*4*.*1*^*-COX8::RFP*^*4*.*2*^*-BFP*_*nols*_^*4*.*3*^*-mCD8::GFP*^*4*.*4*^*-iRFP*_*nols*_^*4*.*5*^*-COX8::RFP*^*4*.*6*^*-mCD8::GFP*^*4*.*8*^*-BFP*_*nols*_^*4*.*9*^*-iRFP*_*nols*_^*4*.*10*^*-BFP*_*nols*_^*4*.*12*^ flies (herein referred to as *PXGS_fluorophores*) as our control lines (**Figure 4a**).

**Figure 4.**
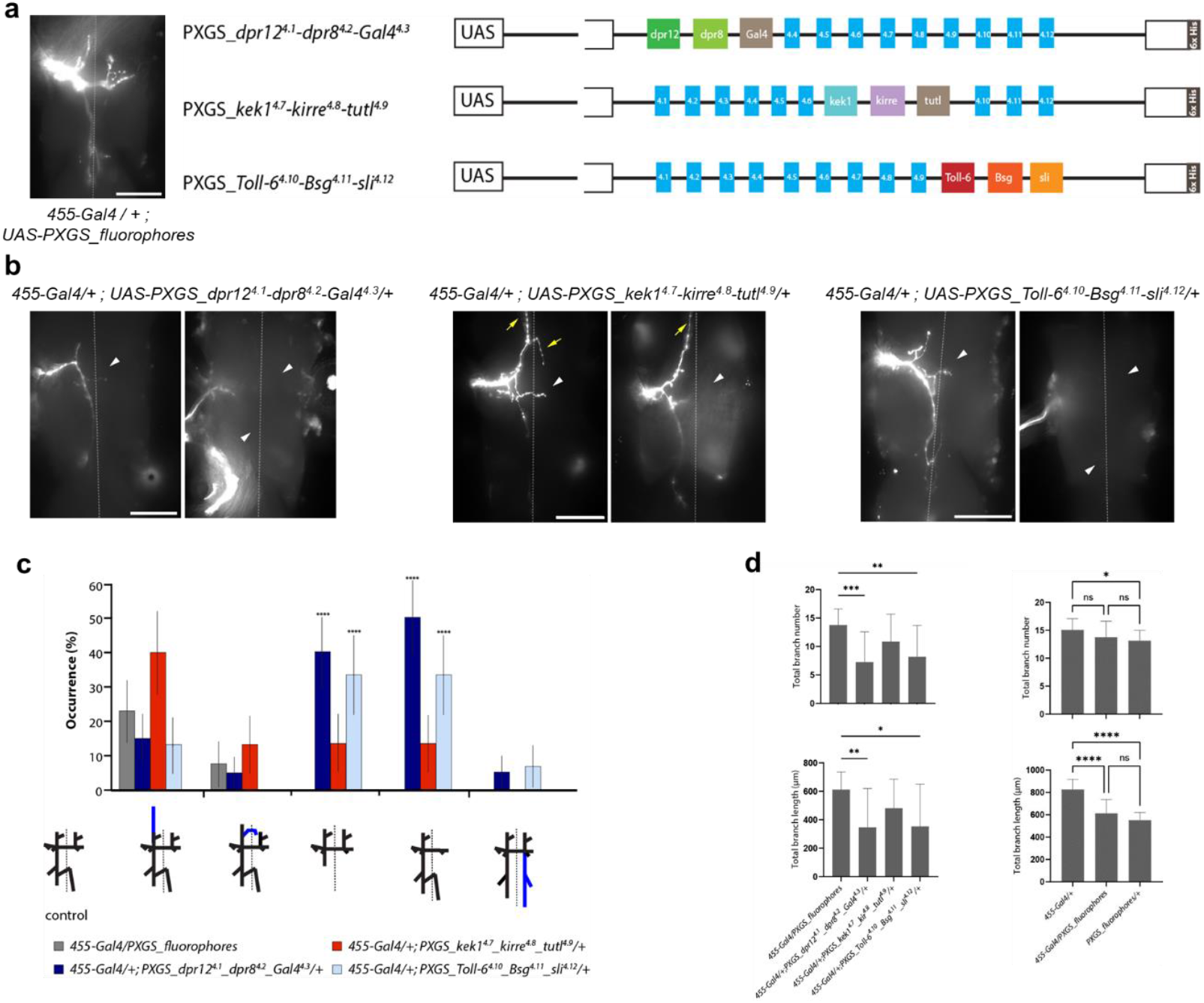
PXGS can express multiple cell surface receptors *in vivo* to manipulate neural wiring. **a**, Representative example of the pSc axonal branching pattern in *455-Gal4; PXGS_fluorophores* controls. A schematic of the three PXGS constructs expressing different combinations of cell surface receptors is shown on the right. **b**, Two representative examples of the pSc axonal branching in *455-Gal4/+; PXGS_dpr12*^*4*.*1*^*-dpr8*^*4*.*2*^*-Gal4*^*4*.*3*^*/+* flies (left images), *455-Gal4/+; PXGS_kek1*^*4*.*7*^*-kirre*^*4*.*8*^*-tutl*^*4*.*9*^*/+* flies (middle images), and *455-Gal4/+; PXGS_Toll-6*^*4*.*10*^*-Bsg*^*4*.*11*^*-sli*^*4*.*12*^*/+* flies (right images). Yellow arrows point to the pSc axon exiting the thoracic ganglion anteriorly. White arrowheads denote missing branches. **c**, Overexpression of multiple cell surface receptors significantly increases axonal targeting errors. A frequency distribution shows the five error types found in the PXGS-cell surface receptor flies. A schematic for each error is shown below the x-axis, where blue branches indicate ectopic branches. Posterior shortening and contralateral anterior missing branches occurred significantly more frequently in *455-Gal4/+; PXGS_dpr12*^*4*.*1*^*-dpr8*^*4*.*2*^*-Gal4*^*4*.*3*^*/+* flies and *455-Gal4/+; PXGS_Toll-6*^*4*.*10*^*-Bsg*^*4*.*11*^*-sli*^*4*.*12*^*/+* flies. **d**, Both *455-Gal4/+; PXGS_dpr12*^*4*.*1*^*-dpr8*^*4*.*2*^*-Gal4*^*4*.*3*^*/+* flies and *455-Gal4/+; PXGS_Toll-6*^*4*.*10*^*-Bsg*^*4*.*11*^*-sli*^*4*.*12*^*/+* flies had significantly reduced total branch length and branch number compared to the control (left graphs). The *455-Gal4/PXGS_fluorophores* and *PXGS_fluorophores/+* both had significantly reduced total branch length compared to *455-Gal4/+*, most likely due to genetic background. *455-Gal4/PXGS_fluorophores* and *PXGS_fluorophores/+* flies had no significant differences in branch length (right graphs). Dotted lines mark the midline. Scale bars are 50µm. Statistical significance comparisons to control are indicated directly above the bar representing each genotype. No asterisks indicate the comparison was not significant. * *p* < 0.05, ** *p* < 0.01, *** *p* < 0.001, **** *p* < 0.0001.

Compared to the control flies (*n* = 13), mis-expression of dpr8 and dpr12 (*n* = 20) or Toll-6, Bsg, and sli (*n* = 15) within the pSc resulted in a significant increase in axonal targeting errors (*p* < 0.0001, *t*-test) (**Figures 4b, c**). The axons frequently failed to extend posteriorly or anteriorly on the contralateral side. In extreme cases, the axon would stall at the entry point (**Figure 4b**). This resulted in significantly shorter total branch lengths and total branch numbers in these two fly lines, compared to *455-Gal4 / PXGS-fluorophores* control (*p* < 0.05, ANOVA) (**Figure 4d**). Mis-expression of kek1, kirre, and tutl simultaneously in the pSc neuron (*n* = 15) frequently resulted in the axon exiting the thoracic ganglion (**Figures 4b, c**).

We also found that the control *455-Gal4 / PXGS-fluorophores* flies had significantly shorter total branch lengths of the pSc axon (*p* < 0.05, ANOVA) compared to *455-Gal4 / +* controls (*n* = 30). However, this is likely due to genetic background differences, and not a “sponging” effect of PXGS on Dscam splicing factors, because the *PXGS-fluorophore / +* flies (within no Gal4) (*n* = 15) also had significantly shorter pSc branch lengths than *455-Gal4 / +* flies. The difference in axonal branch lengths between *PXGS-fluorophore / +* flies and *455-Gal4 / PXGS-fluorophores* flies was not significant (**Figure 4d**).

## Discussion

We created the poly-transgene expression system to allow any *Drosophila* biologist to simultaneously express multiple genes under conditional control. Compared to other multi-gene expression strategies, PXGS offers two distinct advantages. First, it can express many more transgenes than what was previously achievable. Each PXGS construct can express 12 transgenes, so that instead of a standard triple-transgenic animal containing three transgenes, a PXGS animal would have 36 transgenes. Second, the molecular biology to create a PXGS is straightforward and easier to work with. For example, using 2A peptide DNA sequences for more than two genes can be technically challenging due to the repetition of the 2A sequences, which can confound the primer design step, the plasmid assembly, and the sequence verification. This difficulty is eliminated in PXGS. The DNA sequences of the intronic regions spanning each of the 12 exon alternates are different enough that molecular assembly primers do not have multiple binding sites. The turnaround time to generate a PXGS construct is around 3–4 weeks.

Whether a specific cell expresses all 12 Exon 4 alternates for PXGS is dependent on three things. First, the expression level of the Gal4 driver dictates how many rounds of PXGS transcription will occur. This can be supplemented by including the gene for Gal4 within one of the alternates (**Figure 4a**). Second, if the cell expresses *Dscam*, then it should express Exon 4 splicing factors, but all of the splicing factors involved are not known. If *Dscam* exon 4 splicing is deterministic in that cell, then it may never splice all 12 Exon 4 alternates. However, cells that do not express the correct splice factors could be made to, by expressing the correct splicing factors using PXGS itself. Obviously this is dependent on knowing all of the splice factors and mechanisms involved in Exon 4. Additionally, mis-expressing *Dscam* splicing factors would interfere with the endogenous *Dscam* splicing function and may adversely affect the cell given that those Exon 4 alternates were specifically not spliced. Third and similarly, if *Dscam* Exon 4 splicing is random, there still may be insufficient splicing factors despite a high transcription rate. A high transcription rate of PXGS may cause one Exon 4 alternate to be repeatedly selected due to the constitutive recruitment or high local concentration of its splicing factor(s).

For each round of transcription, the probability that any given Exon 4 alternate is not spliced is 11 out of 12. The number of endogenous *Dscam* isoforms expressed within a single cell can be used as a minimum number of rounds of transcription that must have occurred, and this is approximately 10 – 1000 isoforms (Schmucker and Chen, 2009). For ten rounds of transcription, the probability that a specific Exon 4 alternate is not spliced is (11/12)^10^, or 42%. After 50 rounds of transcription, the probability that a specific exon 4 alternate is not spliced is (11/12)^50^, or 1%.

Removing variants in Exon 4 does not affect splicing (Dong et al., 2023). Thus, PXGS can be customized to exactly the number of variants needed, by removing or duplicating the Exon 4 alternates, mimicking what has occurred through evolution (Brites et al., 2013; Lee et al., 2010). If *Dscam* Exon 4 splicing is random, then modifying the number of alternates in PXGS is less likely to negatively affect the cell.

For systems level biological investigations, loss-of-function experiments on multiple genes simultaneously may be required. RNAi is a common approach used for loss-of-function analyses (Qiao et al., 2018). UAS-based expression of dsRNA allows for RNAi-mediated knockdown in specific cells. We generated a *UAS-PXGS_dsRNA-RFP*^*4*.*1*^*-BFP*_*nols*_^*4*.*2*^*-Gal4*^*4*.*3*^*-Shibire*_*ts*_^*4*.*6*^*-iRFP*^*4*.*11*^*-RFP*^*4*.*12*^ fly to knockdown the RFP expression in position 4.12, while leaving the BFP_nols_ in position 4.2 unaffected. Crossing this fly to *nSyb-Gal4*, we found that RFP was not expressed in the brain, but nuclear BFP was clearly visible (**Supplemental Figure 4**). The strong BFP signal was likely due to the Gal4 in position 4.3 that created a positive feedback loop, but this may have also increased the off-target effects of the RNAi, as no iRFP was observed. These experiments demonstrate the potential to use PXGS for polycistronic gene knockdown. Thus, other RNAs may be properly spliced and processed within PXGS, such as guideRNAs, circular RNA, complex secondary structures, or other non-coding RNAs.

The PXGS system will also allow *Drosophila melanogaster* to be a new model system synthetic biology, and for biosynthesis and biomanufacturing in *S2* cells (Coker et al., 2022). For example, we have used PXGS to express the 13 genes required for synthesis of the carbon fixing enzyme, Ribulose-1,5-bisphosphate carboxylase/oxygenase, RuBisCO (Aigner et al., 2017; Gleizer et al., 2019). RuBisCO is famously known as the most abundant enzyme on Earth, because it is expressed at high levels in all photosynthetic organisms. RuBisCO fixes carbon from atmospheric carbon dioxide to convert it into fuel, such as glucose. We crossed the fly *UAS-PXGS_ rbcL1-PQR-raf1*^*4*.*1*^*-cpn60a1*^*4*.*2*^*-cpn60b1*^*4*.*3*^*-rbCS3B*^*4*.*4*^*-cpn20*^*4*.*5*^*-raf2*^*4*.*6*^*-rbcx2*^*4*.*7*^*-bsd2*^*4*.*8*^*-RiBi-Is*^*4*.*9*^*-PRK*^*4*.*10*^*-PGLP1*^*4*.*11*^*-GOX1*^*4*.*12*^ with *nSyb-Gal4* and verified that all 13 genes for RuBisCO were transcribed (**Supplemental Figure 5**). The two genes, *rbcL1* and *raf1* were inserted into position 4.1 with a CHYSEL *Protein Quantitation Reporter (PQR)* linker specific to *Drosophila* (Lo et al., 2015), demonstrating that a poly-cistronic RNA within the poly-cistronic PXGS can generate even larger numbers of transgenes. Thus, PXGS can be used to express complex enzymatic or molecular pathways can in *Drosophila*. All animals in the Tetraconata (Pancrustacea) clade, which is nearly all arthropods, have a hypervariable *Dscam* gene with mutually exclusive splicing (Brites et al., 2013; Lee et al., 2010; Schmucker and Chen, 2009). The PXGS technology can be adapted to any insect or crustacean to express multiple transgenes such as in Sf9 cells from the *Spodoptera frugiperda* fall armyworm, or in High Five cells from the cabbage looper *Trichoplusia ni*, for recombinant protein production *in vitro*, or in mosquitos, silkworm moth, honeybees, and aquaculture shrimp such as the king prawn.

## Methods

### Inserting the Poly-transgene Expression System (PXGS) into plasmids

For simplicity, we refer to *Dscam1* as *Dscam* throughout, because the *Dscam1* gene is the only one of the four *Drosophila melanogaster Dscam* gene paralogs that has the hypervariable mutually exclusive alternative splicing. We amplified the entire *Dscam* exon 4 region along with 300 basepairs (bp) at the 3’ end of *Dscam* exon 3 and 30 bp at the 5’ start of exon 5 from *Drosophila melanogaster* genomic DNA, totaling 6.7 kilobasepairs. The PCR was carried out for 1 min at 94°C, 1 min at 55°C, and 15 min at 72°C for 35 cycles using CloneAmp (Takara Bio Inc. USA, Mountain View, CA) (Celotto and Graveley, 2001; Haussmann et al., 2019; Kreahling and Graveley, 2005). There are several ATG codons (coding for Methionine) within exon 3, which would then create several unintended translation start sites, adding up to 70 unwanted, in-frame amino acids attached to the N-terminus of every PXGS transgene. To eliminate this problem, three point-mutations were inserted into exon 3 (**Supplementary Table 1**).

The *Dscam* fragment was then inserted into the pJFRC7-20XUAS-IVS-mCD8::GFP expression vector (Pfeiffer et al., 2010) using Gibson assembly (Gibson et al., 2009). The fragment was inserted downstream of the UAS sequences and replaced the mCD8::GFP sequence. A 6×Histidine tag was added at the end of exon 5 to distinguish PXGS sequences in reverse transcriptase PCR. Gibson assembly and transformation were performed with the InFusion® HD Cloning Kit and Stellar™ Competent Cells following the kit instructions (Takara Bio Inc. USA, Mountain View, CA).

### Inserting transgenes into PXGS

Transgene insertions into the exon 4 alternates within our different *PXGS* plasmids were created using the Molecular Assembly feature in GeneDig.org (Suciu et al., 2015), primarily using gene synthesis, Gibson assembly, and restriction enzyme digest and ligation. Exon 4 variants were replaced with fluorophore genes codon optimized for *Drosophila*, or cell surface molecule genes, or hairpin RNA sequences.

The *GFP* gene was based on the *mNeonGreen* gene (Ceolin et al., 2020). The *BFP* and *RFP* genes were obtained from a plasmid previously generated in our lab (Lo and Chen, 2019). The *iRFP* gene was based on *miRFP670* (Shcherbakova et al., 2016) and codon optimized for *Drosophila* and synthesized by Integrated DNA Technology (Coralville, IA). To sequester the fluorophores into subcellular compartments to help distinguish their expression, we used transmembrane CD8 (mCD8) for membrane localization, a nucleolar localization signal (nols) for nuclear labelling, and COX8 for mitochondrial localization.

Sequences for the eight cell surface receptors, *Bsg* (798 bp), *dpr8* (891 bp), *dpr12* (1,035 bp), *kek1* (2,643 bp), *kirre* (2,871 bp), *sli* (4,443 bp), *Toll-6* (4,545 bp), and *tutl* (2,892 bp) were obtained from RT-PCR on wildtype flies using oligo d(T) reverse primer to generate the cDNA, and gene-specific forward and reverse primer to amplify the coding sequences. All DNA constructs were verified by colony PCR and Sanger sequencing. All other transgenes used in PXGS were based on their amino acid sequence and then codon optimized for *Drosophila* and synthesized by Integrated DNA Technology (Coralville, IA).

### Cell culture

All PXGS transgenes were first verified using RT-PCR after transfection into *S2* cells. *Drosophila S2* cells were cultured at 25°C in Ex-Cell 420 Medium (Sigma-Aldrich, St. Louis, MO). Cells were transfected with 5 µg of plasmid DNA in 35 mm dishes using Lipofectamine (ThermoFisher, Waltham). 48 hours after transfection, RNA was extracted from the S2 cells and reverse transcription was performed on transfected and untransfected cells. PXGS-specific reverse primers for the 6×His tag or for the overlap between exon 5 and the vector were used to identify PXGS transgenes, and oligo-d(T) primers were used for endogenous *Dscam* mRNA alone. Endogenous *Dscam* exon 4 was used as the positive controls, and PXGS plasmid with no *Actin5C-Gal4* transfection was used as the negative controls. PCR was performed using gene-specific forward and reverse primers.

### Fly stocks

To reduce variability due to environmental conditions and genetic background, flies were reared at 25°C on standard cornmeal under 12h light / 12h dark cycles. To express PXGS fluorophores in different cell types, the pan-neuronal *nSyb-Gal4*, glial *Repo-Gal4*, ubiquitous *Tubulin-Gal4*, and scutellar *455-Gal4* drivers were used. *PXGS* plasmids that were verified for expression in *S2* cells were used to make transgenic animals. Transgenic *Drosophila* lines were generated using PhiC31 integrase-mediated transgenesis performed by BestGene, Inc (Chino Hills, CA).

### Carbocyanine dye labeling and Drosophila imaging sample preparation

*Drosophila* mechanosensory axonal arbors were labelled using carbocyanine dye injection onto the pSc neurons, as previously described (Chen et al., 2006; Cvetkovska et al., 2013; Dos Santos et al., 2019; Kays et al., 2014; Neufeld et al., 2011). The left and right pSc bristles of 2-day old female flies were plucked and the thorax was left overnight in fixative solution (3.7% paraformaldehyde in 0.2 M carbonate-bicarbonate buffer, pH 9.5). The left and right pSc neurons were labelled using the carbocyanine tracers, DiI or DiD (ThermoFisher). The flies were left undisturbed and partially immersed in 0.2 M carbonate-bicarbonate buffer at pH 9.5. After 48 hours, the thoracic ganglion was dissected and imaged.

### Image acquisition

Fluorescent images were acquired using an AxioScope A1 epifluorescence microscope (Carl Zeiss) with a 40× objective (N.A. 1.0) or an FV1000 laser-scanning confocal microscope (Olympus) with a 40× objective (N.A. 1.3). For mechanosensory axon imaging, 10–20 images were taken at different focal planes. Maximal intensity z-projections were stacked to generate the final image. Images were adjusted only for contrast and brightness when required. Transmitted light images were used to evaluate dissection and dye labeling quality and to measure thoracic ganglia width.

### Image analysis

Only images without central nervous system damage or surface occlusions were included for data analysis. Measurements of axonal branch length and branch number were performed blind to genotype by randomly distributing the axonal arbor data of the total dataset. Custom-written software in MATLAB was used to perform the quantitative image analysis.

To determine the baseline variability in wildtype pSc mechanosensory neurons, the total number of branches and branch lengths per arbor were measured from thirty control 455-Gal4 / + animals. A skeleton of the control 455-Gal4 / + axonal arbor was then created based on the frequency of occurrence of each branch, which was found to be consistent with our previous measurements (Chen et al., 2006; Cvetkovska et al., 2013; Dos Santos et al., 2019; Kays et al., 2014; Neufeld et al., 2011). We defined an ectopic branch as any branch that has less than 30% occurrence in the wildtype. Representative examples of axonal arbors in figures were randomly chosen from the sets of all axonal arbor images of the genotype.

### Statistical analysis

Statistical significance for total axonal branch length and number of branches was determined using one-way ANOVA followed by Dunnett’s multiple comparisons test. Statistical significance for frequency of occurrence was determined using a two-tailed *t*-test for proportions. Statistical analysis was performed using GraphPad Prism version 8.0.0 for Windows (GraphPad Software, San Diego, California USA) and SPSS (version 25). The datasets, analyses, and materials used in the current study are available from the corresponding author upon request.

## Acknowledgments

The authors thank Farida Emran for assistance with experiments. This work was supported by a grant from the Natural Sciences and Engineering Research Council of Canada (2020-04902, B.E.C.). The funding agency had no role in the design of the study and collection, analysis, and interpretation of data and in writing the manuscript.

## Author Contributions

B.E.C. designed the experiments and supervised the project. R.Y.Y. and A.B-M. performed experiments and analyzed the data. R.Y.Y., A.B-M., and B.E.C. wrote the manuscript.

## Competing Interest Statement

R.Y.Y., A.B-M., and B.E.C. are inventors on a pending patent describing the system and materials in this manuscript.

The data that support the findings of this study are available from the corresponding author upon request. Correspondence and requests for materials should be addressed to.

Supplementary Information is provided as five supplementary figures and one supplementary table.

