## Supplementary Information for "PXGS: a Poly-Transgene Expression System based on Mutually Exclusive Splicing of *Dscam*"

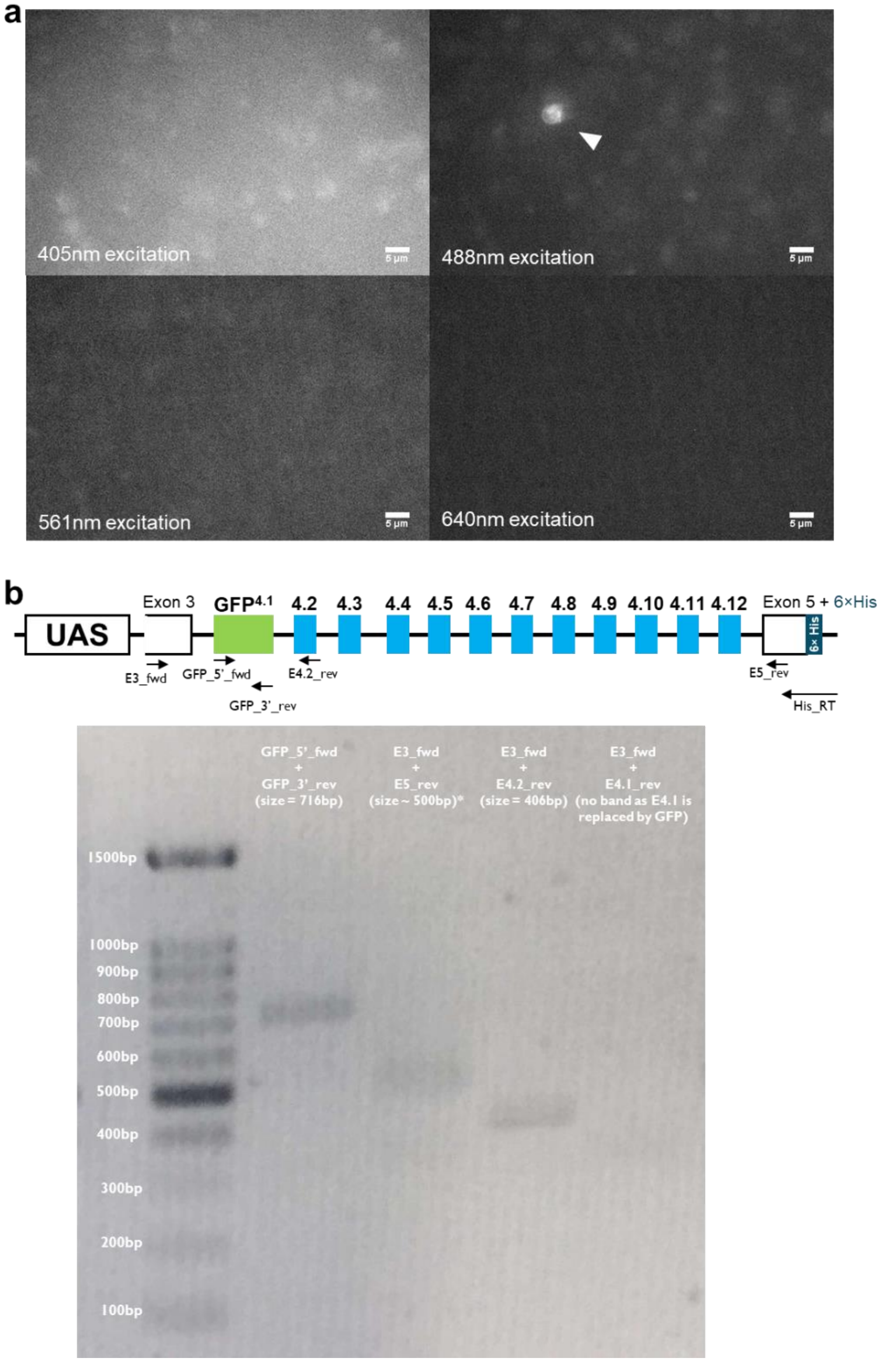


**Supplemental Figure 1** – Complete replacement of an Exon 4 alternate can properly splice and produce functional protein. **a,** Replacing Exon 4.1 with the gene for *GFP* produces green fluorescence when co-transfected with *Actin5C-Gal4* in *S2* cells. Blue (upper left), green (upper right), red (lower left), and far red (lower right) channels are shown. Scale bars are 5µm. **b,** PXGS-specific RT-PCR of different exon 4 variants in transfected *S2* cells verified their appropriate splicing and expression. *GFP* mRNA band is expected at 716 bp, the remaining possible exon 4 variants are expected at ~500 bp (denoted with a *), Exon 4.2 is expected at 406 bp, and Exon 4.1 is not expected.

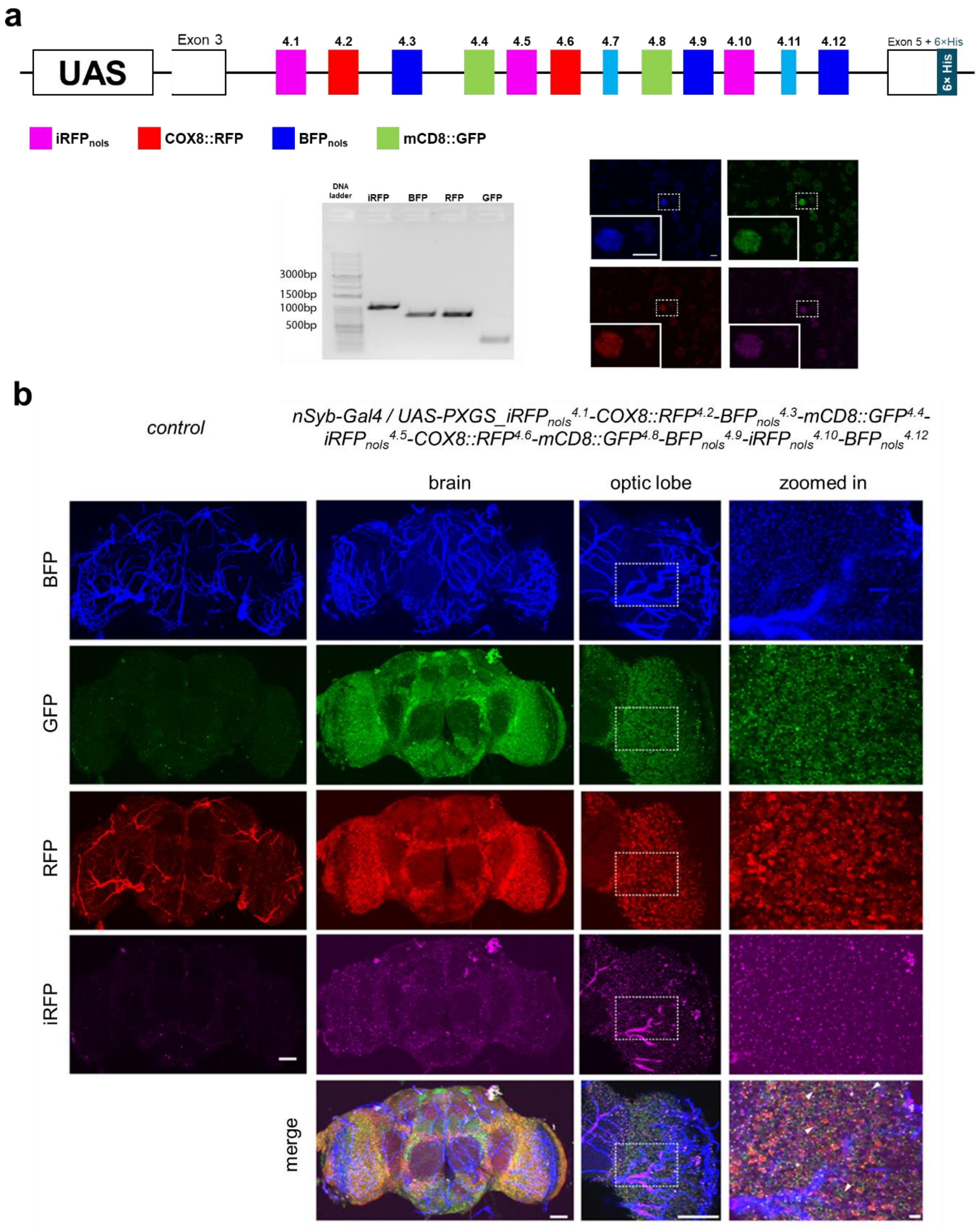


**Supplemental Figure 2** – Replacing 10 of the 12 Exon 4 alternates with fluorescent protein transgenes increases the expression pattern in neurons. **a,** Co-transfection of *UAS-PXGS_iRFP*$_{nols}^{4.1}$-*COX8::RFP*$^{4.2}$-*BFP*$_{nols}^{4.3}$-*mCD8::GFP*$^{4.4}$-*iRFP*$_{nols}^{4.5}$-*COX8::RFP*$^{4.6}$-*mCD8::GFP*$^{4.8}$-*BFP*$_{nols}^{4.9}$-*iRFP*$_{nols}^{4.10}$-*BFP*$_{nols}^{4.12}$ fly with *Actin5C-Gal4* in *S2* cells produced mRNA at the expected sizes (*iRFP* = 1044bp, *BFP* = 810bp, *RFP* = 804bp, *GFP* = 270bp), and produced fluorescent cells. Scale bars are 10µm. **b**, Crossing the pan-neuronal *nSyb-Gal4* driver line with the transgenic fly *UAS-PXGS_iRFP*$_{nols}^{4.1}$*-COX8::RFP*$^{4.2}$*-BFP*$_{nols}^{4.3}$*-mCD8::GFP*$^{4.4}$*-iRFP*$_{nols}^{4.5}$*-COX8::RFP*$^{4.6}$*-mCD8::GFP*$^{4.8}$*-BFP*$_{nols}^{4.9}$*-iRFP*$_{nols}^{4.10}$*-BFP*$_{nols}^{4.12}$ resulted in all neurons expressing all four fluorophores at higher levels than the four fluorophore PXGS fly. The differential subcellular localizations of the fluorophores can be observed at higher magnifications in the optic lobe. White arrowheads in the zoomed in merged image (bottom right) point to cells that express all four fluorophores evenly. Scale bars are 50µm, 50µm, and 5µm respectively from left to right.

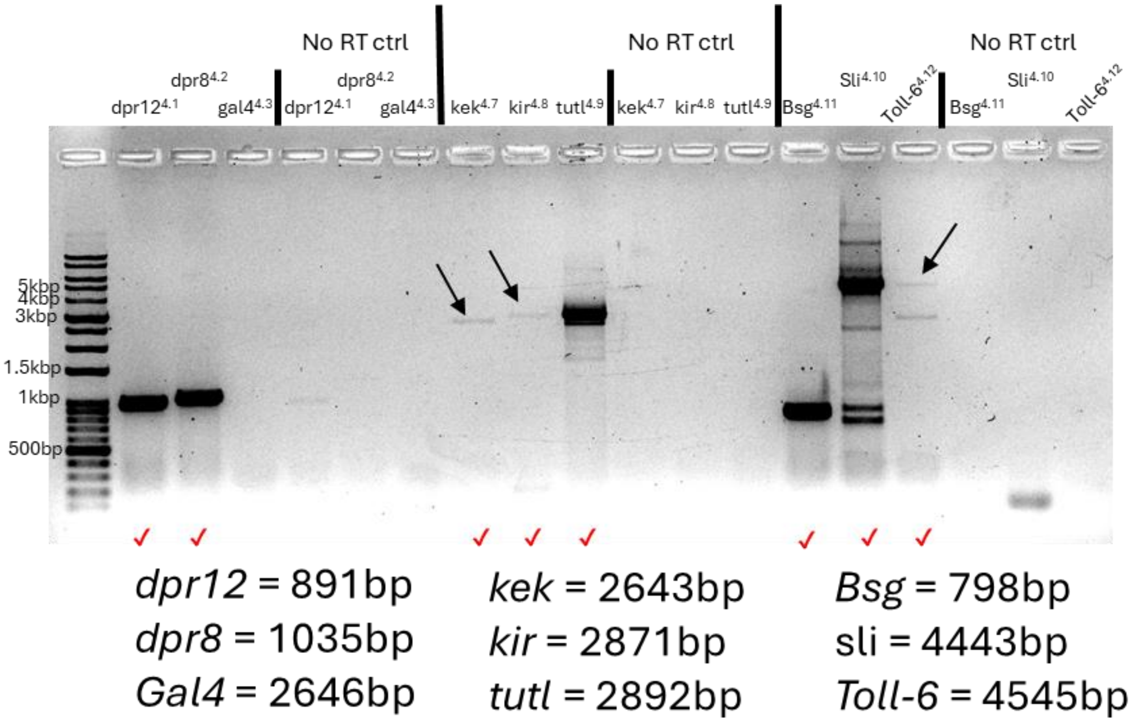


**Supplemental Figure 3** – PXGS can express multiple cell surface receptors in *S2* cells. *UAS-PXGS_dpr8*$^{4.1}$*-dpr12*$^{4.2}$*-Gal4*$^{4.3}$, *UAS-PXGS_kek1*$^{4.7}$*-kirre*$^{4.8}$*-tutl*$^{4.9}$, and *UAS-PXGS_ Toll-6*$^{4.10}$*-Bsg*$^{4.11}$*_sli*$^{4.12}$ were each co-transfected into *S2* cells with *Actin5C-Gal4*. RT-PCR on each gene was performed 48 hours after transfection. The PCR products for *UAS-PXGS_dpr8*$^{4.1}$*-dpr12*$^{4.2}$*-Gal4*$^{4.3}$ are shown in the left side, *UAS-PXGS_kek1*$^{4.7}$*-kirre*$^{4.8}$*-tutl*$^{4.9}$ in the middle, and *UAS-PXGS_ Toll-6*$^{4.10}$*-Bsg*$^{4.11}$*_sli*$^{4.12}$ on the right. The expected sizes for each gene's PCR product is shown. *Gal4* mRNA from PXGS expression was not detected likely due to a mutation within the upstream intronic sequence that may interfere with its splicing.

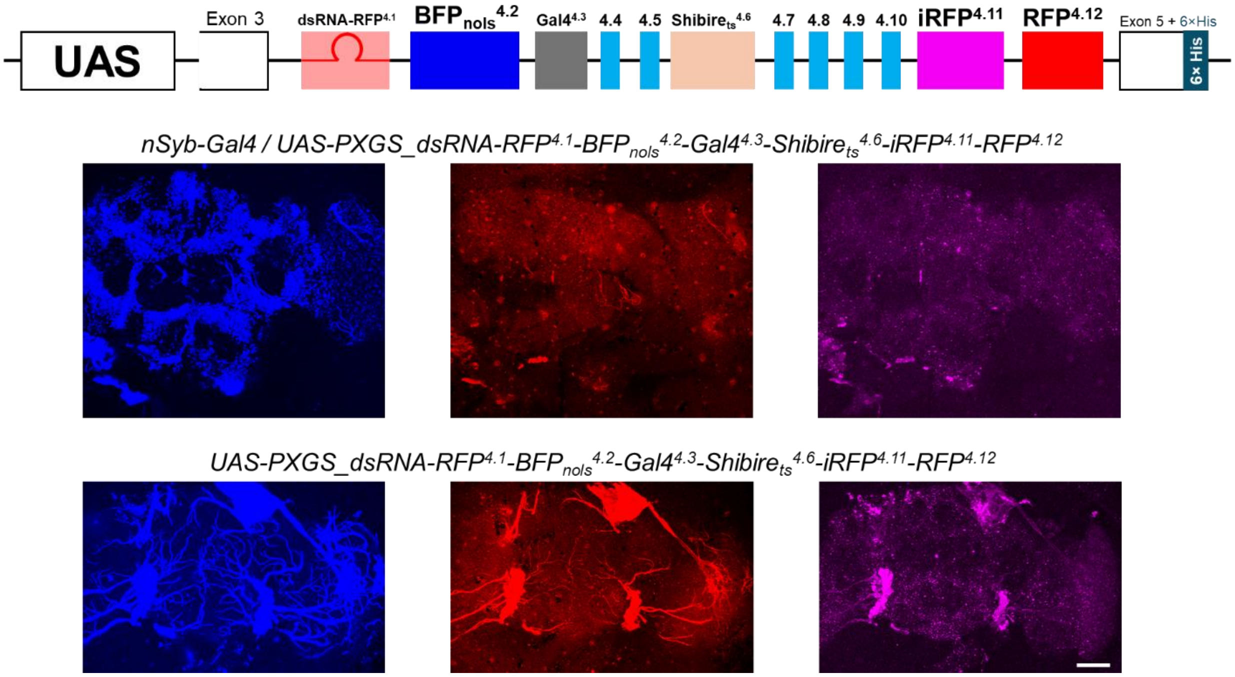


**Supplemental Figure 4** – PXGS can express dsRNA for RNAi knockdown. Flies expressing dsRNA against RFP did not have red fluorescence, but still expressed BFP. The low iRFP signal is possible due to the off-target effects of the RNAi. Scale bar is 50µm.

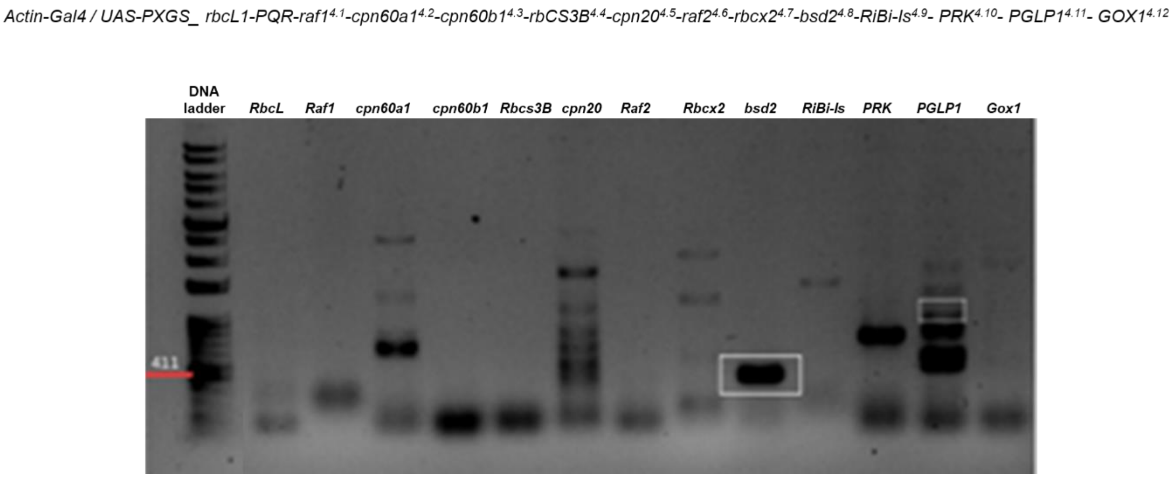


**Supplemental Figure 5** – PXGS can express 13 large genes to re-create synthetic biology pathways. Actin-Gal4 flies were crossed to PXGS flies expressing 13 genes to synthesize the carbon fixing enzyme, Ribulose-1,5-bisphosphate carboxylase/oxygenase (RuBisCO). RT-PCR verified that all 13 genes were expressed in the flies.

**PXGS_empty_annotated**

Exon3

Exon 3 mutations:

A -> **t**: get rid of start

T -> **a**: create a stop

A -> **t**: get rid of start

Exon 4.1 to Exon 4.12

Exon 5

His tag

TGCCGACCAAAAAGGACCCGTCTTCCTCAAGGAACCCACCAACC
GCATTGACTTCTCCAACTCCACGGGCGCAGAGATCGAGTGCAAG
GCCAGCGGCAATCCC**t**TGCCCGAGATTATTTGGATCAGGAGCGA
CGGTACCGCCGTGGGTGATGTGCCCGGATTGCGTCAGATCTCAT
CCGACGGCAAGCTGGTCTTCCCTCCATTCCGCGCCGAGGACTA
CCGCCAGGAGGTCCATGCCCAGGTGTACGCCTGCCT**a**GCCCG
CAACCAGTTCGGATCCATTATCTCCCGGGACGTCC**t**TGTGCGAGC
CGGTAAGTTGTCAGGGTTCAGAGGTAAACCAGGGGCAGTGCAAA
GTAAATGTTGGTAATTGACAAAGAAAGTCATTCTTAAAAAAGGATAGA
TATGAGCTTTATATATTTTTAAAAAGTTTAAAAAAATATTTAATAATAATAT
ATAAAATATTACAAATATTCATTTAAATGAAACTCAACATTTAGTTTAAC
AGATTTTTAATACAAATCTGCAATCAAAATACAAGAATAGAAGATGAA
AAAGCTGCGAGTTAAAGTAACTAATTATTTTGAAGTCGGTAGAAAATT
CCTTTTGAATTAGGCAACCAACAGGTGACTACTTTGCACTGCCCTG
AGTTTACCTTTCTTTTACAGGTTCACCCAAATGAAGGCCTATGTATCT
GCGTTGGTTTCGTATCTCCACTACTTGGCACTTTCAGTTTGCAGCTA
AAAGTTGCCGGTTGCCTTTGCGTGTCAAGTCGCTCTCTGCAGTCTT
TAACCCCTTGTGTTTGACTTATTGTTGCACCGGCTGCGGCAATTAAA
ACCGAGTTTTCACGGGATTTCCATCGAACGCTCTGCGAATTGCCA
AAGTTTTCCGCACGCTTTCATTTTCACACATTTCTTAGTCGAACTATG
CAACGTTTAGAGGAGTTTAATGCAAGTTGGCATTTCAGTTTAATCGG
TTCGTTTTCGCAGCTTTACTTCCTGTCGCAATGTTTTGTCGGCAACT
GTACCTGTCATGCCCGTGCATGTCTGTTCAGCAACCTTTAACAGCT
TCCCTGGAAGATGTGGCAGCTGCGACGATGACGATGATGAAGAAG
ATGATGAAACTGTCAGACTATTTAGCGCACACCGTGGGGTTGACG
GGTTGCCAGCTGATCGAGTTATGGACCAGACCCAACTCCCTTGGA
TATCAGTCACTTCCGCTTCCATGGAGCCATCCCTAAATGGGCCAG
ACTTGAGCACATTTTAGCTAATTATGTCGGAGCCATGCCACCCGCT
TACTCATCGCCAGCTGACGTGTTCCCTGGCAATTAGCCAGCACAC
CTATCCCAGACACCACCTCTCGACCCCCGTTCCAGAAGCCTGAC
ATCATTATTGTCGTTTGCTTTCGCTTGCTATTAACCGGAGGAATTTAT
CATTTTATAGTATGCTTATGTGTGAATAAGGCGCAGACGAGGAAAAC
ATTCTGCACAGCCGGCCCGCAGCTAGTTGGCCGAAATGAGTTATG
GTCTACACCCTGAACACGGGCTGGCGTAAAGAATATTTTGGTTAATT
GAAATTAACTTTTGCCTGTCAAGCAGCGGCGCGATGATAGTTTCGG
GGTTTTCGACCAGAAGGTCGTTTACAACGGCAGAATTTTCCAGGAA
CTCAGTTCCTAAGCACTTTCAAGGTTGTGAGAATGTTTTGGACATAAT

| | |
|---|---|
| | AGGTAATAATCGGCCTTTTCCCAGTGGTTGCCCAGTACTACGAGG<br>CGGATGTTAACAAGGAGCACGTTATAAGAGGCAATTCGGCGGTCA<br>TCAAGTGTCTGATTCCATCCTTCGTGGCCGATTTCGTCGAAGTGGT<br>GTCCTGGCACACCGATGAGGAGGAGAACTACTTTCCGGGCGCG<br>GAATACGGTGCGATCCGAGCTGCAGATGCAGGCAGATAGGATAC<br>ATGTGTAGTTTTGCTCTACTAAAAGCATGCTGGGGTCGCGATTCCG<br>CTAGTTGCACAGAAAATCCCTGCATTCCATCATTCATTCATTCCTTAT<br>TTTAATCTGAATCTCACCACTATTATCGTTTCTCCTACCTGTTTAGTCG<br>TTTCCCAACACTACGAAGAAGATATACACAAGGCATTTGTCATCCG<br>CGGCAATTCCGCCATATTGAAATGCGATATACCGTCGTTTGTAGCC<br>GATTTCGTGAATGTTATATCCTGGCATAGTGACGAGAAAGAGAATTT<br>CTATCCAGGCACTGAATATGGTGTGTGACTAGAAGAACAGTCTCTGT<br>TTGTGAGCTTTTAGAAAACTATCCTTAAACTACCTTAACCGCTACTGT<br>ACAACACTGCAAAAAAGGCAAACTCACTGAAACACTTGTGCACAG<br>TGCAAGGGCAGTTAACGGGAAACACTAATTACAAAATTCTGTAAAAT<br>ATACAACACACCAAACTCAAAGAATAACGTATTAACCAACACTGGC<br>TAGGCAGAAACCAGTCGAGAATCCTTCGCTCCCTCGTCGTAGCTT<br>TCCCAATTCGTCAGAGTTTCGTTTTCAGTTCTACCCAGAATTTGATTT<br>CCTTTCGGTTTTATTACATAAACGCATTTATTTGATTGCTGTCATTGCT<br>GTTTTAGTCGTTGCCCAATACTACGACACCGATGTAAATAAGGCCTA<br>TGTAATACGCGGCAATGCGGCGGTTCTGAAATGCGAAATCCCCTC<br>CTTTGTGGCCGATTTTGTGAGCGTTGTCTCCTGGCACACCGATCAG<br>AACGAGGACTTTCTGCCAGGCAGCGAATACGGTTTGTTGGCCAAC<br>TTTTTGATTACATTTTGGGCTAGCTATACATACATAGGGTTTGTACAGT<br>TGCGTGCCACCTAAGAATTATGGATAAGTACCTAACCATTTTCCCTC<br>CTGCATCTATTATGTTTACTTTTATGCGTTTATCTTCGCTATCTCGGGT<br>GACTGCGTGTTGATGAGCCATGTTATCAAAACCATTATTAACCTCTT<br>GCCCGGAGATTTTCTCAGGGAAGCGCGGAGCTCGCGAGCTGATT<br>GCGCTTAGGGTCCCATTAAGTGCACTCCGCGATATTGTCTGCGCT<br>GTTAGAAGACACTCTCGTTTCAATTAGCTAGCTTCACATTTTCACAC<br>GCCCGCCGCACGCAAGGCTTGGACTTAAACTTGGGTTCTGTAGG<br>CGCCCCATCCCCCATCTTTCCCTCCGTACCATCGCATCATTATGTA<br>ATCAGTGGCTATGTGAACTCACCTTCAGTTGTGCAGCAGTTCTATGA<br>ATCGGAGGTCAACAACGAGTACGTCATAAGGGGCAATGCAGCGG<br>TGCTGAAGTGCTCGATTCCATCCTTTGTTGCGGACTTTGTGCAGGT<br>CGTATCCTGGCAGGACGAAGAGGGTCAGCTCTACGGCTCGCTAG<br>GCGATCAACAGGGAACGGGTACACATGCACTATAGCGAAACCTAA<br>TGGTTATGTAGGATTATCCACAGCCCATCCCGCACCTACCCCAAT<br>CCTTCATTCAGCAATTTCATTTCATTTTGGTTCCTACTCATAAAAACTC<br>TTGCTTTACAGTCGTGAATCAGTATTACGAGGCTGAGGTTGTTTCCG<br>AGTACGTTATCCGTGGCAATACGGCTGTATTGAAGTGCACTATTCC |

| | |
|---|---|
| | CAGTTTTGTAGCCGACTTCCTGCGGGTCGAGGCCTGGGTGGCCA<br>GTGACGGCACTGAGTACGCTCCGGAAGAGGATTTTGGTACAGTCC<br>CGCACCGTTGAAACTATTAAGAAGTTAGTCCTAAGCAGCAATCCCA<br>TTCCTTCCCCATTTCAATTTGCTTCAGCTTCTGTTTCATTTCACCACT<br>CTACCCCATTTTAAATCGCAGTTGTGAATCAGTTTTACGGCGCCGAT<br>ATCCTGATGGAGTATGTCATCAGGGGAAATGCGGCGGTTTTGAAAT<br>GTTCCATTCCGTCGTTCGTGGCCGATTTTGTGCGTGTCGAGTCCTG<br>GATAGATGAGGAAGGCACCGAACTGCGTCCCTCTGAGAACTATGG<br>TATTACCGCAGAGTCAAAAATTCTTGCAACGGCAACACTTTGTGGG<br>GGTTTCGTTCCTATCCCATCCTATCTCCAACTTTTCTATATCTTTCCGT<br>TATTGCACACCTTTGCAGTTGTCAATCAGTTCTATGAGGCGGAGATT<br>ATGACCGAGTATGTTATCAGGGGTAATGCTGCCGTTCTGAAGTGTTC<br>CATCCCGTCGTTCGTGGCTGATTTTGTTCGAGTTGAGTCCTGGATC<br>GATGATGAGGGCAATGTCTTGTCCTTTTCGGACAATTACGGTGCGA<br>GGCAAAGATGTTCATCCAGTAGTTTTATGCTTAATTTCCACCAGCTC<br>CATTCCCAGTGATCCCCTCTTTTCCATTTACTTATTATTTTATTCGATT<br>CAAAGTCGTTTCACAGTTCTACATAACCGAGGCCGAGAATGAGTAC<br>GTAATTAAAGGGAACGCGGCGGTGGTCAAGTGTAAGATCCCATCC<br>TTCGTGGCCGATTTCGTGCAGGTTGAGGCCTGGGTGGACGAGGA<br>GGGCATGGAGTTGTGGCGCAACAATGCCACCGACGCCTATGGTA<br>CAGTTCAAGCAAAACATACTGCTTCGGTTAAGTGCAGCACTAGGTC<br>CCAACAACCACTTCCTTGACTCCCAACCCCCAAACCCATATTTTA<br>CACTACATTATTCCTTTGTTTTTCTATCGACTCAGTGGTCATCCAAAG<br>CTATGAATCGGAGGCGGACAATGAGTACGTCATTAGGGGCAACTC<br>TGTGGTGATGAAGTGCGAGATTCCCTCCTACGTGGCCGACTTCGT<br>GTTCGTCGATCTCTGGCTGGACTCGGAGGGTCGCAACTACTATCC<br>GAACAATGCCGCAGAGACGGGTACTTGGCCAGGTTTCTTTGCCGT<br>AGCTCTCTCAGGCTCCTCTATAGCTATCCAGAAACCCATTCCCGTT<br>CACTTCGGTTATTATTTCACCAAGTTCGCTCTGATTTCCTTCAGTTGT<br>CCACCAGTTCTACCAGACACGGGTCATCGATGAGTTTGTGCTGCG<br>TGGCAACTCCGCCACCTTGAAGTGCTTGGTGCCCTCGTTTGTGGC<br>AGACTTCATCGATGTCGAGGGTTGGATCGACGAGGAGGGCGTGG<br>AGATCCTGCGGCCCCTCCCGGACGACTCCGTTGGTAATCACTAC<br>GACTTTCGGCATTAGCATAGTTAGAGTACGAAACCTTCAACCCACC<br>GACCCTTCCCTGTCCCGATTTTTAGTTTCTCACCTCCCGTCTTGCA<br>GTGGTGAAGCAGTTTTTCGAGTCGCAAGTCTACGACGAGTATGTGA<br>TCAAGGGCAATGCGGCCATTTTCAAGTGCCAGACGCCCTCCTTTG<br>TGGCGGATCACATAGACATAACCGACTGGATCGATACCGAGGGC<br>GAGGTCTTCACGAAGAACATAACTGGTAGGAGTCCGAGTCTCGTA<br>GATTTGTGCCGTTTTCTCAGTTACATGAATGGGGATCCTCTTTCAGA<br>CGCCTCTTTTATCCTTATGATTTAACCCCACGTCCCACTCACTCAAT |

| | |
|---|---|
| | CAATGTCTTCGTACACTTTGCAGTTGTGCCACAATCATACACCGTCA<br>ATGTCATGGACGAGTCCATACTGCGAGGCAATAGCGCCATCCTAA<br>AGTGCCACATTCCCAGTTTTGTGGCCGATTTCATAGTGGTCGATTC<br>GTGGGTGGAGGACGAGGAGCGAGTTATCTATCCCCAGGAGGATAT<br>CGCGGAAAGCGGTAAATTTACAGACTAAGAGCTCTCGCATTGCCC<br>ACTCCTCCGTAACATCATCCTGCGTCCCCTATCCAATTCGGTTCGC<br>TTCGATCTGGTCTCGATGTGGTTTCTTTAGAGCTCTGTAAAAGAAGTT<br>CTCCACGTTCAGCGCTGCTTAGGCTTTTGAATGGCCTTCATTGTGT<br>ATTTCTTTCATTCTGTCCGCCTTTTCTTTCTATGTTTTTTGTTGTTTTTTTT<br>TGTTTTCCCATCCCAACCCTTTTTAGTTTTCCCCTCCAAAAATCCAT<br>GGGTCGCTCCTGACTTGCCCAAAAAGAAGCCGAAACCCCTTCAT<br>GTAGTTGAACGTAGCGTTGATAAAGTAGCGTAGTGTAAATAAATGCA<br>AAAATGCTAATAACCCATCGACCACGTAACAATATGTAAATACATATC<br>ATTTCAAACGAATAATCCAACATTGCTGTAATTGTATATACCATAAAGA<br>TTCGAGTATATATAATATGTAACATGCATTTCATCTAAGATTATTGGCAT<br>TTTTGTTTCATGCATGTCTAGAATTAAGCACAATGCCATGGAATTTGG<br>AGAATCACGATCGAAGCAATCAACTAAAACTATTTGAATTGTAAAAAT<br>GTATCTCAATTCGAAATTTAAATAGATTACAATCAATCTTTCCGGTTTG<br>ATTCTGTTTGTTTATACTACAAACTCTTGGGCATTGTTGGTGTATAGTA<br>AATGTTTATGGAACTTTGTTAAAACAAATTCGATTGAAATCAAAATCAA<br>AGACTAAAGGAGATCCTAACAAGCTTAGTAGTTAATCTTGCTCTAAA<br>AAATTTCATAAATGAATCAAATTAATAACGAATTCTATTTTTAATCAAAC<br>ATGTGCGAAATAACTACAAGGGCTTCTCATTTAGTCTATAGATCTTAA<br>CTTATCCTTAATCATTTCAAAGTCACATTGCATGGTCAACGATTAGCA<br>GCATTCAATTTTTGCACAATTAAAAGTAACACAAAATGAAAAATGATT<br>ACCAGCCATGTGGCTTTCAAATGTAGAGCGAATTAATTCAATTGAAT<br>GCATGCTAGTCAATTGATTTTGTTCGCCGTACTGTATTGTTAATAAGTT<br>AAGTTAGCCGACAATTCATCACTAACATATTCTTTATTCTCTATCTGCA<br>CATATCAAAATATCAGATGGAAAGTACCTGGTATTGCCCTCTGGAGA<br>GCTGCACATCCGTCATCATCACCATCACCACTAA |

**Supplemental Table 1** – DNA sequence of PXGS construct with mutated Exon 3, endogenous Exon 4, and 6×Histidine tagged Exon 5.
